# Complement C3-targeted gene therapy restricts onset and progression of neurodegeneration in chronic mouse glaucoma

**DOI:** 10.1101/369181

**Authors:** Alejandra Bosco, Sarah R Anderson, Kevin T Breen, Cesar O Romero, Michael R Steele, Vince A Chiodo, Sanford L Boye, William W Hauswirth, Stephen Tomlinson, Monica L Vetter

## Abstract

Dysregulation of the complement system is implicated in neurodegeneration, including human and animal glaucoma. Optic nerve and retinal damage in glaucoma is preceded by local complement upregulation and activation, but whether targeting this early innate immune response could have therapeutic benefit remains undefined. Because complement signals through three pathways that intersect at complement C3 activation, here we targeted this step to restore complement balance in the glaucomatous retina, and to determine its contribution to degeneration onset and/or progression. To achieve this, we combined adeno-associated viral retinal gene therapy with the targeted C3 inhibitor CR2-Crry. We show that intravitreal injection of AAV2.CR2-Crry produced sustained Crry overexpression in the retina, and reduced deposition of the activation product complement C3d on retinal ganglion cells and the inner retina of DBA/2J mice. This resulted in neuroprotection of retinal ganglion cell axons and somata despite continued intraocular pressure elevation, suggesting a direct restriction of neurodegeneration onset and progression, and significant delay to terminal disease stages. Our study uncovers a damaging effect of complement C3 or downstream complement activation in glaucoma and establishes AAV2.CR2-Crry as a viable therapeutic strategy to target pathogenic C3-mediated complement activation in the glaucomatous retina.

## INTRODUCTION

Glaucoma irreversibly impairs vision by progressive neurodegeneration of retinal ganglion cells (RGCs). Axonopathy precedes neuronal loss, with the earliest damage detectable in unmyelinated axons at the optic nerve head (ONH), followed by gradual axonal and neuronal decline and dropout with advancing age^1^. The pathobiology of this neurodegenerative disease remains unclear due to manifold processes and risk factors, including age, ethnic group and elevated intraocular pressure (IOP)^2^. Current therapies can successfully lower IOP and decelerate but not halt the progression of neurodegeneration^3^. Thus, developing neuroprotective approaches to interrupt progression is a primary goal^4^. To be effective, therapies should target pathways and processes that have been demonstrated to drive neurodegeneration in animal models and human glaucoma.

Emerging research ties glaucomatous neurodegeneration to complement dysregulation in the retina and ONH, identifying C1q and C3 as candidate therapeutic targets^5, 6^. Complement is an innate immune molecular network that acts to remove pathogens, damaged cells and immune complexes^7^. C3 is the central complement component and point of convergence for three main initiation pathways (classical, alternative, and lectin)^8^. C1q is the pattern recognition molecule that canonically initiates the classical pathway, but has broader functions beyond the complement system^9^. In the CNS, C1q and C3 tag synapses, as well as damaged and apoptotic neurons for clearance by microglia or infiltrating macrophages^10^. Neurons, astrocytes, microglia and peripheral monocytes are local sources of complement proteins, especially during damage or disease^11^. Thus, in addition to traditional roles in serum immune surveillance, complement is now recognized for its central roles in CNS development and adult homeostasis, as well as in pathology, including aging and neurodegeneration^12, 13^. How local CNS complement contributes protective, reparative, or pathogenic roles is under active study.

In the retina, changes in complement expression underlie aging and a number of pathologies, including glaucoma^14^. Retinal proteome and transcriptome analyses have detected C1q and C3 upregulation in human glaucoma^15, 16^ ^17^, and ocular hypertension^18, 19^, as well as across animal models of induced IOP elevation^5, 15, 19-27^. Expression of C1q mRNA was detected in RGCs and ONH microglia/macrophages^25, 27, 28^. These findings place local complement dysregulation during early stages of glaucoma, preceding optic nerve (ON) degeneration.

The DBA/2J mouse strain develops progressive RGC degeneration with variable age of onset exhibiting disease features relevant to human glaucoma^29^. Systemic knockout of C1q and C3 genes in the DBA/2J was the first attempt to assess the impact of complement activation in glaucoma. C1q deficiency was neuroprotective, delaying but not halting progression of ON degeneration^27, 28, 30^. Unpredictably, C3 deficiency worsened degeneration before the terminal stage^26^. Both IOP elevation and anterior segment disease were delayed by C1q, but not C3, deficiency. Of note, DBA/2J mice are deficient for C5 and terminal pathway activation, implying that upstream complement components are sufficient to drive neurodegeneration, although restoring C5 worsens its severity^31^. These divergent outcomes of C1q and C3 deficiency suggest that there may be activation of multiple complement pathways during the course of chronic glaucoma, each functioning with partial independence, possibly at different stages and contributing pathogenic or protective roles. Since gene knockout cannot distinguish systemic from local effects, developmental from adult effects, or onset versus progression of neurodegeneration, the therapeutic potential of modulating complement activation in the glaucomatous retina remains undefined.

Inhibitors affecting the central step of formation of C3 convertase formation and cleavage/activation have become a focus for development of therapeutics that effectively modulate complement activation, regardless of the initiation pathway^32^. Furthermore, the deposition of C3 activation fragments, iC3b, C3dg and C3d, covalently bound to cell surfaces, serves as a long-lived indicator of local complement activation amenable to imaging in the living animal or *ex vivo*, and as ligand for CR2-fused inhibitors to achieve localized regulation of complement activation^8, 33, 34^. In rodents, the main regulator of C3 convertase formation is Crry, a structural and functional ortholog of human complement receptor 1 (CR1)^35, 36^. Endogenous Crry is expressed on brain microglia and astrocytes^37^, as well as across the retina and RPE^38^, and upregulated in the DBA/2J retina and ONH^27^. Site-targeted CR2-Crry, which directs soluble Crry to activated C3 deposits^39^, inhibits local complement activation 10-fold more actively than untargeted soluble Crry^33^.

Endogenous complement inhibitors have long been used in C3-targeted strategies to interrupt initiating complement pathways and broadly balance complement activation^32^. To overcome adverse systemic effects and effectively inhibit local C3 activation, targeted approaches have been designed to selectively supply complement regulators to sites of cleaved C3 deposition on cells and tissues^33^. CR2-Crry is a fusion protein of the murine C3 convertase inhibitor Crry (complement receptor-1 related protein Y) and the C3d/dg complement receptor 2 (CR2) moiety, which targets Crry to sites of activated C3 deposition^39^. CR2-Crry has been proven neuroprotective in the ischemic brain^39-41^, as well as in the experimental autoimmune encephalomyelitis model of multiple sclerosis^42^. Systemically administered CR2-Crry crosses defective blood-brain barriers in these models of acute CNS injury^34, 43^, but would have limited capacity to cross an intact barrier.

Ocular gene therapy viral vectors, such as adeno-associated virus (AAV), have been used to provide the diseased retina with local and permanent expression of Crry and other inhibitors, to chronically attenuate complement activation in models of age related macular degeneration^44-46^. Serotype 2 AAV vectors yield widespread transduction of the inner retina and RGCs 1 month after intravitreal injection in the mouse eye^47^, and over 10 months in the DBA/2J retina^48-51^. Thus, stable expression of targeted complement inhibitors via AAV gene therapy should be a viable approach to chronically attenuate complement activation during DBA/2J glaucoma progression.

Here, we use targeted gene therapy to test whether C3 activation contributes to the onset and/or late progression of neurodegeneration in glaucoma by using AAV2 to deliver CR2-Crry. DBA/2J mice received intravitreal injections of AAV2.CR2-Crry at 7 months of age, when most ONs are structurally intact, and were aged and evaluated for alterations in timing and severity of neurodegeneration relative to AAV2.GFP and naïve controls. We provide evidence that AAV2.CR2-Crry retinal gene therapy effectively increases Crry expression and dampens C3d deposition in RGCs, which results in significant long-term neuroprotection of ONs and retina. Degeneration was almost eliminated at 10 months, and progression to terminal stage was suppressed at 12 and 15 months. Thus, AAV2.CR2-Crry ocular gene therapy provides a precise and translatable strategy to locally balance retinal C3 activation during disease progression and reduce neurodegeneration in chronic glaucoma.

## RESULTS

### Intravitreal AAV2.CR2-Crry limits the deposition of C3d on RGCs

To allow local expression of targeted inhibitors of C3 activation to the retina of DBA/2J mice, we adopted ocular gene therapy using high-efficiency triple Y-F mutant capsid AAV2 vectors, which result in widespread and stable transduction of RGCs and inner retina in adult mice^47^. We performed bilateral, intravitreal injections of AAV2.GFP reporter at 10^10^ total vector genomes per eye in 7 month-old DBA/2J mice, and confirmed efficient transduction of the inner retina in 10 month-old mice, consistent with previous reports of DBA/2J viral gene therapy^48-51^. Retinal wholemounts and radial sections showed GFP expression in the nerve fiber layer (NFL) and ganglion cell layer (GCL) (**Figure 1A, B**). GFP expression was also observed in interneurons and Müller glia (**Figure 1B**), but was not detectable in photoreceptors, microglia, vasculature, or the retinal pigment epithelium (RPE). Coimmunostaining with the RGC-specific transcription factor Brn3a and use of Thy1^+/CFP^ DBA/2J reporter mice^52^ confirmed AAV2.GFP expression in RGCs and their axons across all quadrants and eccentricities (data not shown). The ON showed GFP-expression along axons within the proximal segment (**Figure 1C**). This spread of AAV2.GFP transduction in the DBA/2J retina and optic nerve is consistent with previous reports of AAV2-GFP transduction, which also show its presence in the lateral geniculate nucleus and superior colliculus^53^. AAV2.GFP expression in the retina was first detected 2 to 3 weeks post-injection and peaked by 1 month, with expression persisting for at least 5 months (data not shown). To selectively reduce complement activation in the glaucomatous retina the original construct for recombinant CR2-Crry fusion cDNA^39^ was packaged in YF-mutant capsid AAV2 vector^47, 54^, then 7-month-old DBA/2J mice received single injections of AAV2.CR2-Crry bilaterally, and were aged to 10 months together with naïve littermates. To detect overexpression of Crry, whole retina qPCR was performed to measure the relative fold change of total Crry (endogenous plus exogenous) and to compare treated to naïve retinas. Expression of Crry mRNA showed an 8-fold mean increase in AAV2-CR2-Crry-treated retinas (*n*=8) over naïve (*n*=4) and AAV2-GFP (*n*=8), with 10 to 30-fold upregulation in 6 out of 8 treated retinas (**Figure 1D**). One-way ANOVA revealed a significant difference between the three groups (*p*<0.001). Levels of Crry in AAV2.CR2-Crry were significantly elevated from naïve and from AAV2.GFP controls by a Tukey’s multiple comparisons test (*p*<0.01 and *p*=0.0011, respectively), whereas AAV2.GFP controls were not significantly different from the naïve condition (**Table 1**). Due to differences in variances between samples, we confirmed these results with an unpaired Student’s t-test with Welch’s correction.

**Figure 1.**
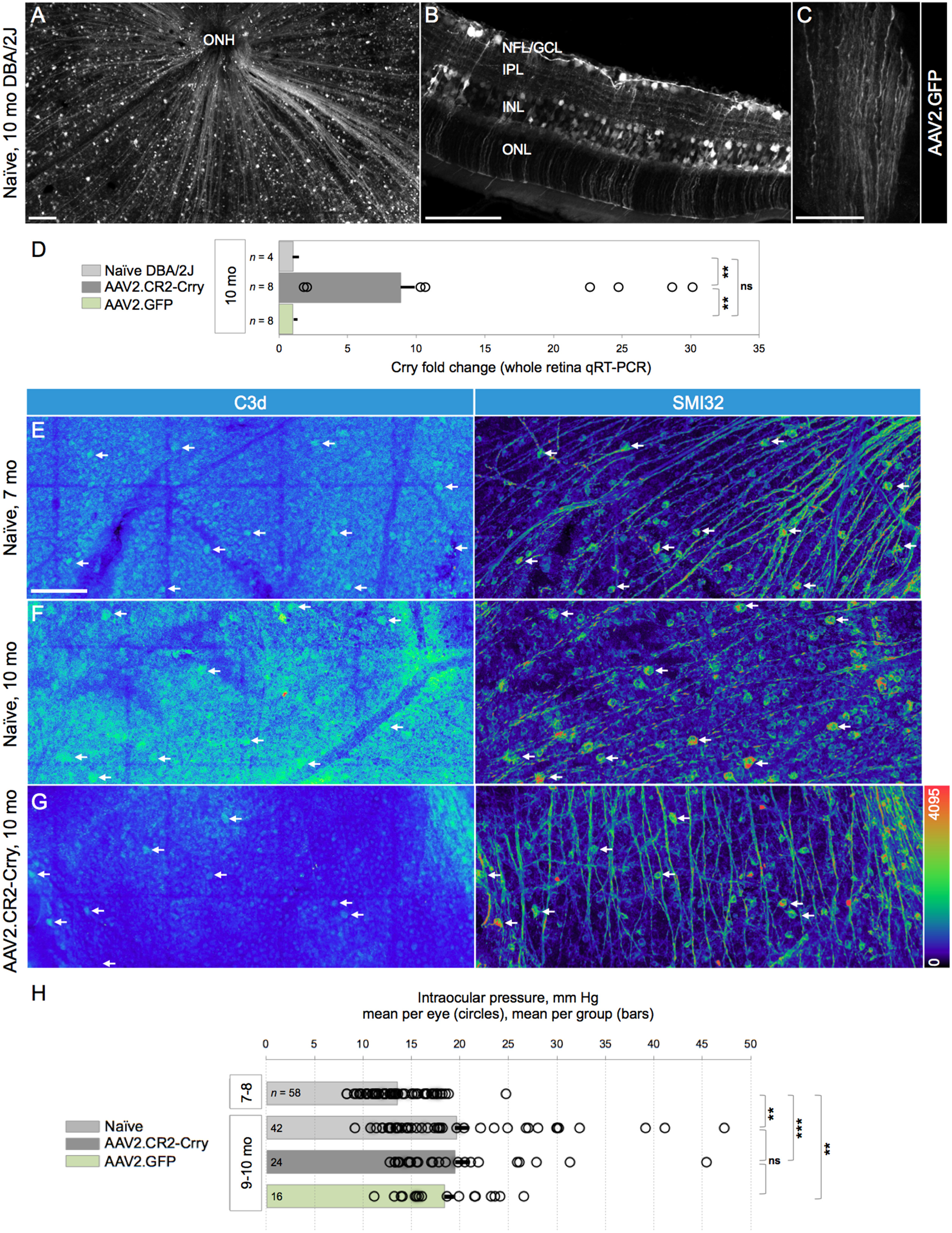
C3d deposition on RGCs and GCL is lessened by retinal Crry overexpression. (**A**) Representative retinal wholemount of 10 month-old (mo) DBA/2J mice injected with AAV2-GFP at 7mo, showing widespread GFP localization to RGCs and their axons. Confocal image of the NFL and GCL, showing the innermost 25 μm below the vitreal surface, seen as maximal-intensity projection, and low magnification. (**B**) Radial section across the retina, displaying GFP expression in RGC somata and axons, IPL interneurons, and Müller glia. Confocal image of a 10 μm-maximal intensity projection. (**C**) ON showing GFP content in axons along a longitudinal cryosection, viewed in a single slice (0.8 μm-thick), ~2 mm behind the retina. (**D**) Mean fold change of Crry mRNA expression in whole retinas from 10 month-old mice treated with AAV2.CR2-Crry or AAV2.GFP relative to age-matched naïve retinas measured by quantitative RT-PCR. Error bars represent mean ± s.e.m. Dots represent fold change in single retinas. \*\**p*< 0.01 each (see Table 1). (**E-G**) Confocal images of retinal wholemounts spanning the NFL and GCL (~40 μm-maximal intensity projections; mid-periphery), showing single channels in pseudocolors to emphasize variations in signal intensity. (**E**) Naïve retina, representative of 7mo DBA/2J, double-immunostained for C3d and SMI32. C3d localized to GCL cells throughout large sectors (left), and to SMI32+ αRGCs (arrows). (**F**) Naïve retina, representative of 10mo mice, which display intense C3d deposits throughout the GCL, and in SMI32+ αRGCs (arrows). (**G**) AAV2.CR2-Crry-treated retina, representative of 10mo mice, showing C3d immunostaining in the GCL confined to reduced retinal sectors, and to few declining SMI32+ αRGCs (arrows). Scale bars, 100 μm (A, B), 50 μm (C-F). (**H**) Bar graph of IOP (mean ± s.e.m per group) showing that at 9-10 months of age, eyes in all groups underwent a significant increase compared to 7-8 months naïve eyes (*p*<0.01). The IOP of individual eyes (circles) exhibit extensive overlap at 9-10 months, regardless of condition, with means not significantly different between naïve and AAV2.CR2-Crry or AAV2.GFP (see Table 2). Number of eyes per group indicated within each bar. Significance level is indicated as follows: ns (not-significant), * (*p*< 0.05), ** (*p*<0.01), *** (*p*<0.001), for this and following figures.

**Table 1.**
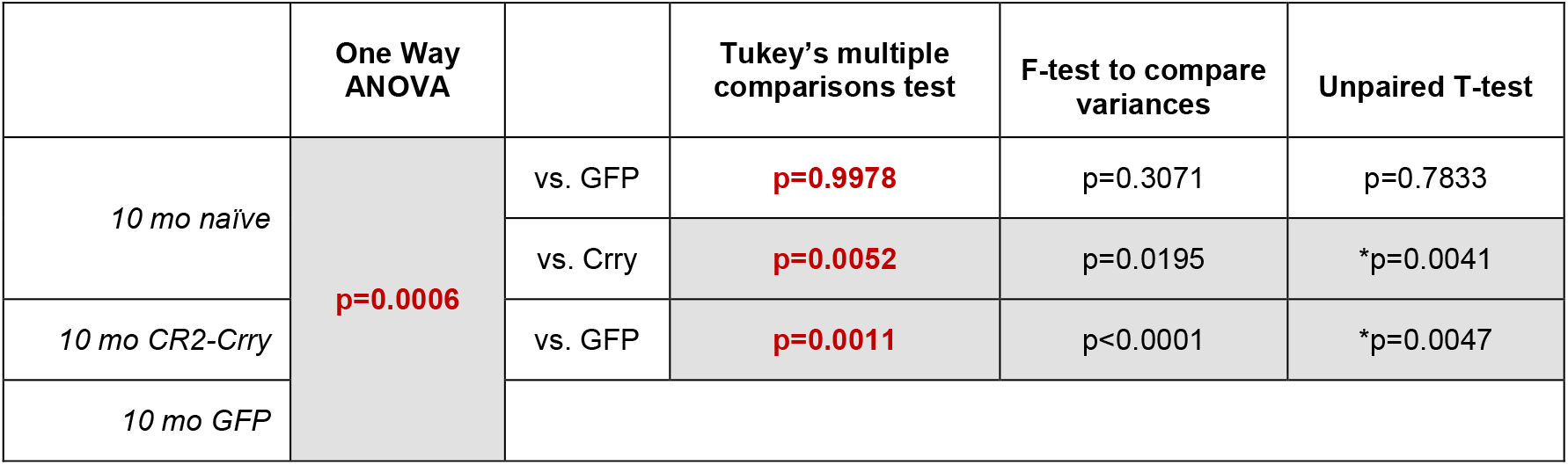
Crry mRNA expression. Statistical comparisons for Delta CT values (Figure 1D). * indicates Welch’s correction. Shaded cells highlight significant differences, and red font identifies the values reported in Results (this applies to this and all tables).

To gauge C3 activation in the retina, we performed immunostaining of retinal wholemounts to detect the C3 breakdown product C3d, which is deposited on cell surfaces^33, 55^. Confocal microscopy of naïve 7 month-old DBA/2J showed C3d staining of cell somata in the GCL, with variable levels of fluorescence intensity across large retinal sectors (**Figure 1E**). This deposition was also detectable by immunostaining in retinas at 5 months of age (data not shown). To confirm labeling of RGCs we performed coimmunostaining of αRGCs with the SMI32 neurofilament antibody^50, 56, 57^, which consistently revealed C3d deposition in αRGCs across multiple eyes (*n*=8; **Figure 1E**). C3d deposition persisted in 10 month-old naïve DBA/2J mice, which showed labeling of the GCL throughout entire retina, with generally higher intensity levels relative to 7-month naïve mice (**Figure 1F**). Use of Thy1^+/CFP^ DBA/2J reporter mice and triple-immunostaining, showed C3d colocalization to Thy1-expressing RGCs, which colabeled with SMI32 and/or gamma-synuclein in retinal wholemounts (**Figure S1A**). The C3d antibody has been used in previous studies^58-60^, and to further test its specificity, we immunostained retinas from non-glaucoma Gpnmb^WT^ DBA/2J mice prepared as radial sections and observed negligible C3d staining, comparable to negative immunostaining controls of naïve DBA/2J retinas omitting primary antibody (data not shown).

Retinas treated with AAV2.CR2-Crry at 7 months exhibited a general reduction in C3d deposition at 10 months relative to naïve retinas at the same age, with retinal wholemounts displaying uniformly low levels of C3d expression throughout the GCL, and fewer cells with high C3d deposition (**Figure 1G**). Dystrophic SMI32-labeled αRGCs were stained for C3d, while most αRGCs with healthy morphology had dim or undetectable C3d staining. We did not observe C3d deposition on microglia of naïve or treated retinas (**Figure S1B**), unlike reports of C3d localization to brain microglia in patients with multiple sclerosis and other neurodegenerative conditions^61-63^. We did detect C3d on GFAP+ parenchymal astrocytes in some naïve retinas aged 10 months (data not shown). Together, these observations suggest that AAV2-CR2-Crry is able to decrease the activation of C3 throughout the retina in DBA/2J mice aged 7 to 10 months.

### AAV2.CR2-Crry retinal gene therapy does not alter the typical elevation of intraocular pressure in DBA/2J

To determine whether the IOP of mice treated with AAV2.CR2-Crry would elevate with similar time course and values as naïve DBA/2J mice, we compared IOPs from 7 to 10 months of age. By a Kruskal-Wallis test, we found a significant difference across all groups (p<0.0001). As expected, IOP increased significantly in naïve mice from 7-8 months (13.52 mm Hg ± 0.42 s.e.m., *n*=58) to 9-10 months (19.6±1.37 mm Hg, *n=*42; *p*<0.01 by Kolmogorov-Smirnov), consistent with published results^64, 65^ (**Figure 1H**). Importantly, we find a comparable and significant IOP elevation from 7-8 to 9-10 months for both AAV2.CR2-Crry-treated (19.48±1.53 mm Hg, *n=*24) and AAV2.GFP-injected eyes (18.40±1.15 mm Hg, *n=*16), (*p*<0.01 each; **Table 2**). Furthermore, there was not statistical difference in the mean IOP between 9-10 month-old naïve, AAV2.CR2-Crry and AAV2.GFP conditions by Kruskal-Wallis comparison (*p*=0.812). These results were confirmed with a post-hoc Dunn’s Multiple Comparisons test (**Table 2**). These findings indicate that intravitreal gene therapy with AAV2.CR2-Crry or AAV2.GFP did not reduce or delay IOP elevation in treated DBA/2J eyes.

**Table 2.**
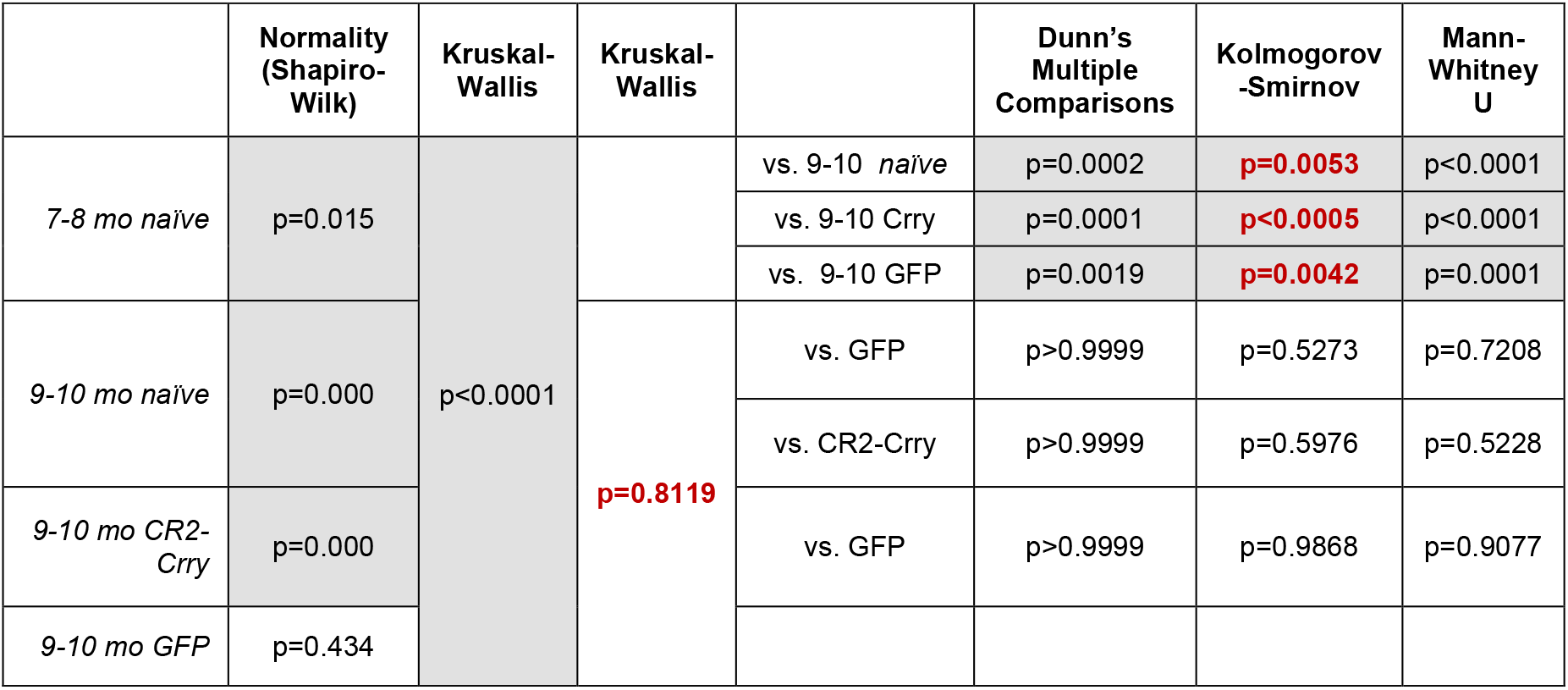
Intraocular pressure. Statistical comparisons for mean IOP per group (Figure 1H).

|  | Normality<br>(Shapiro-Wilk) | Kruskal-Wallis | Kruskal-Wallis |  | Dunn's<br>Multiple<br>Comparisons | Kolmogorov<br>-Smirnov | Mann-Whitney<br>U |
| --- | --- | --- | --- | --- | --- | --- | --- |
| 7-8 mo naïve | p=0.015 | p<0.0001 |  | vs. 9-10 naïve | p=0.0002 | p=0.0053 | p<0.0001 |
|  |  |  |  | vs. 9-10 Crry | p=0.0001 | p<0.0005 | p<0.0001 |
|  |  |  |  | vs. 9-10 GFP | p=0.0019 | p=0.0042 | p=0.0001 |
| 9-10 mo naïve | p=0.000 |  | p=0.8119 | vs. GFP | p>0.9999 | p=0.5273 | p=0.7208 |
|  |  |  |  | vs. CR2-Crry | p>0.9999 | p=0.5976 | p=0.5228 |
| 9-10 mo CR2-Crry | p=0.000 |  |  | vs. GFP | p>0.9999 | p=0.9868 | p=0.9077 |
| 9-10 mo GFP | p=0.434 |  |  |  |  |  |  |

### Complement inhibition reduces RGC degeneration rates and severity

To establish whether the local attenuation of retinal C3 activation could alter the course and severity of RGC somal degeneration and loss, we performed intravitreal injection of AAV2.CR2-Crry in 7 month-old DBA/2J female mice and aged them to 10 months. We then examined their level of RGC preservation *ex vivo*, relative to both age-and gender-matched control AAV2.GFP-treated and naïve eyes. To detect RGCs with persistent transcriptional integrity, retinal wholemounts were immunostained with a Brn3 antibody that recognizes all three isoforms of Brn3 transcription factors^66-68^, since their downregulation serves as an early marker of RGC dysfunction^66-69^. Confocal microscopy images spanning the GCL of entire wholemounts showed a range in the density of Brn3+ RGCs. All experimental groups showed healthy retinas with uniformly dense Brn3+ RGC mosaics (**Figure 2A, D**) and declining retinas with Brn3+ RGC loss across sectors radiating from the ONH across one or more quadrants (**Figure 2B, E**). However, degenerative retinas with prevalent low Brn3+ RGC densities and sectorial depletion were common in naïve and AAV2.GFP control eyes, but infrequent in AAV2.CR2-Crry eyes (**Figure 2C**).

**Figure 2.**
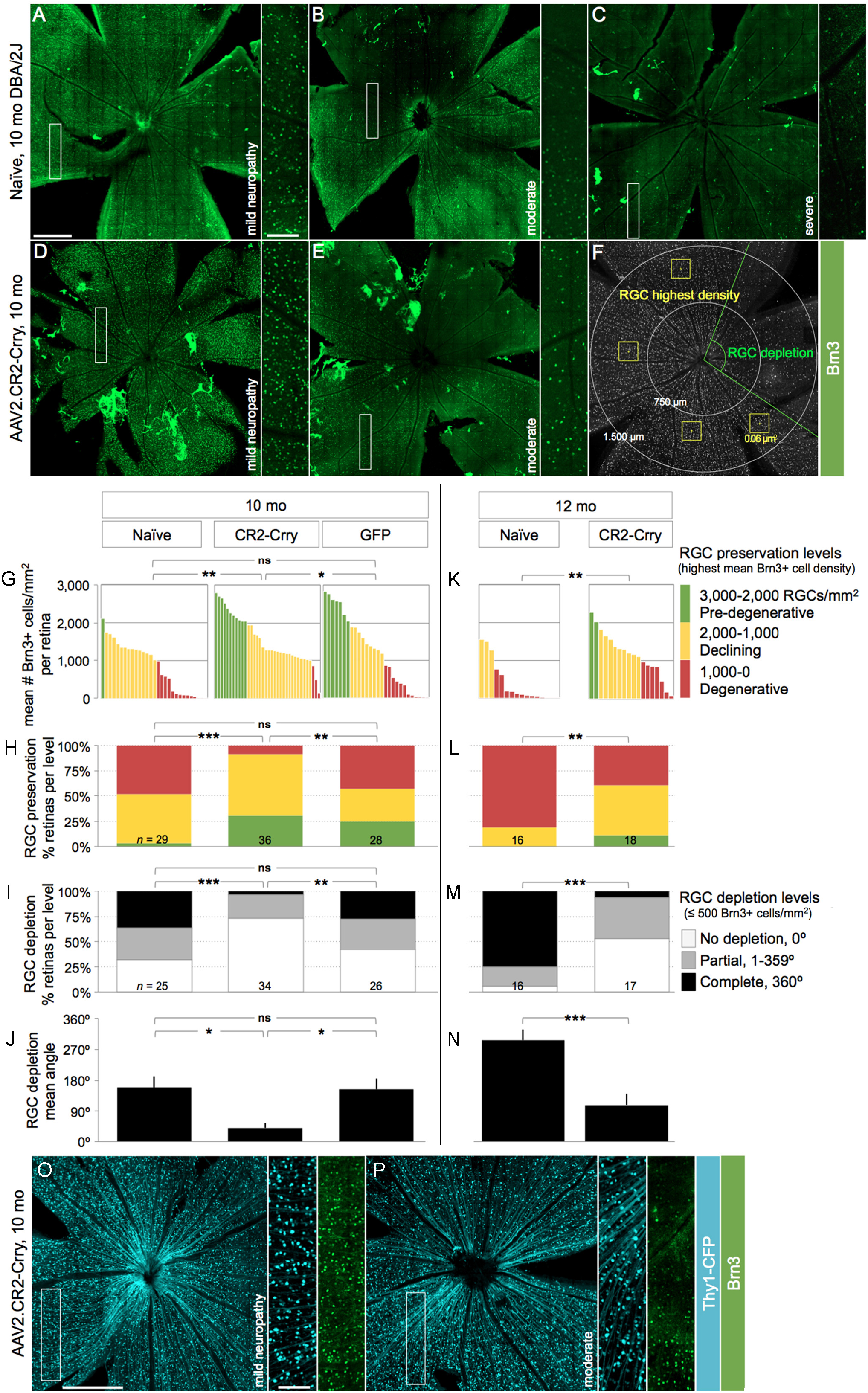
AAV2.CR2-Crry reduces Brn3+ RGC degeneration onset and progression. (**A-E**) Confocal images through the NFL and GCL of retinal wholemounts immunostained for Brn3, shown as maximal intensity projections of 25-35 μm. Insets display RGC mosaics at high magnification. Retinas representative of 10 month-old naïve (**A-C**) and AAV2.CR2-Crry-treated mice (**D and E**), with corresponding level of ON damage indicated as pre-degenerative, moderately or severely degenerative. (**F**) Example of Brn3+ cell sampling method. (**G, K**) Plot of the highest density of Brn3+ RGCs (mean number nuclei/mm^2^) for individual retinas at 10 and 12-months of age respectively. (**H, L**) Stacked bar chart representing percent retinas at each level of RGC preservation per condition: pre-degenerative (mean Brn3+ RGC density/mm^2^ > 2,000, green), declining (1,000-2,000, yellow), and degenerative (< 1,000, red). (**I, M**) Stacked bar chart representing percent retinas at each level of RGC depletion per condition: not depleted (0° with less than 500 Brn3+ cells/mm^2^), partial (1-359°), and complete (360°). (**J, N**) Bar graph of mean depleted retinal area (degrees) per condition. (**O, P**) Confocal images of Thy1^CFP/+^ RGCs from representative retinas of 10 month-old DBA/2J mice. Insets show their Brn3 co-expression at high magnification. Scale bars, 500 μm (A-F, O-P), and 100 μm (insets). See Tables 3-6 for statistical analysis.

To represent the variable patterns and severity of RGCs loss that are characteristic in DBA/2J glaucoma progression^57^, in individual retinas we estimated “RGC preservation” by counting Brn3+ nuclei within the densest sector at the mid-periphery of each quadrant, and also estimated “RGC depletion” by measuring the extent of retinal sectors with ≤ 500 Brn3+ nuclei/mm^2^ (**Figure 2F**). Analysis of mean Brn3+ cell densities revealed a large variability in all experimental conditions at 10 months of age, ranging from 0 to almost 3,000 cells/mm^2^ (**Figure 2G**). We found a significant difference across all 10mo groups by Kruskal-Wallis (*p*<0.01). Due to the variability of the model and therefore, non-normal distribution of preserved Brn3+ cell density, we performed pairwise comparisons of these distributions by non-parametric Kolmogorov-Smirnov tests (**Table 3**). There were significant differences between 10 month-old AAV2.CR2-Crry treated retinas (*n*=36) compared to both naïve (*p*=0.01; *n*=29) and AAV2.GFP retinas (*p*<0.05, *n*=28), but no significant difference between naïve and AAV2.GFP retinas (*p*=0.103). The non-parametric Mann-Whitney *U* test, which is more sensitive to differences in medians, confirmed the significant difference of AAV2.CR2-Crry compared to naïve (*p*<0.001) but found no significant difference compared to the AAV2.GFP group (*p*=0.117; **Table 3**), suggesting a partial effect from AAV2-GFP delivery over naïve retinas.

**Table 3.**
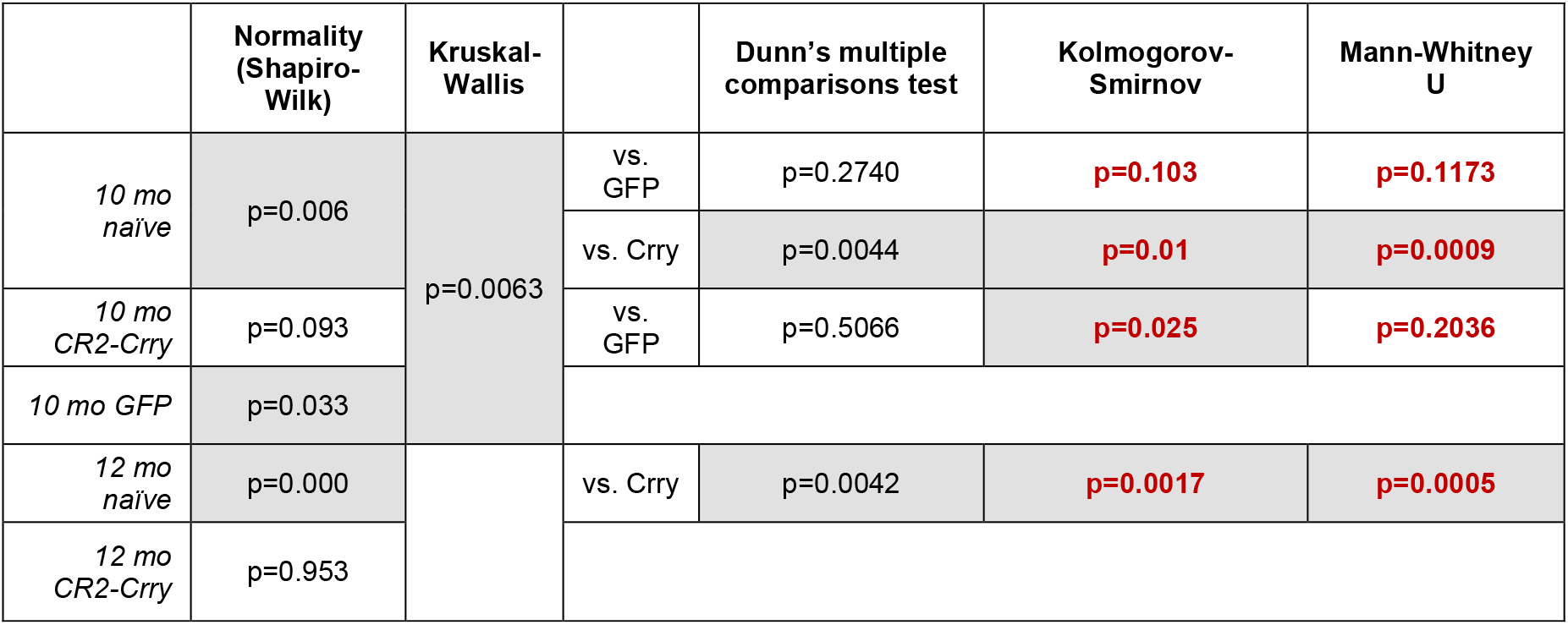
Highest mean Brn3+ cell density. Statistical comparisons for mean number of Brn3+ cells/sq.mm per retina (Figure 2G and K).

|  | Normality<br>(Shapiro-Wilk) | Kruskal-Wallis |  | Dunn's multiple<br>comparisons test | Kolmogorov-Smirnov | Mann-Whitney<br>U |
| --- | --- | --- | --- | --- | --- | --- |
| *10 mo naïve* | p=0.006 | p=0.0063 | vs. GFP | p=0.2740 | **p=0.103** | **p=0.1173** |
|  |  |  | vs. Crry | p=0.0044 | **p=0.01** | **p=0.0009** |
| *10 mo CR2-Crry* | p=0.093 |  | vs. GFP | p=0.5066 | **p=0.025** | **p=0.2036** |
| *10 mo GFP* | p=0.033 |  |  |  |  |  |
| *12 mo naïve* | p=0.000 |  | vs. Crry | p=0.0042 | **p=0.0017** | **p=0.0005** |
| *12 mo CR2-Crry* | p=0.953 |  |  |  |  |  |

To better represent the stages of retinal degeneration per condition, we compared the proportions of retinas at each level of Brn3+ RGC preservation (highest mean density), as pre-degenerative (3,000-2,000 RGCs/mm^2^), declining (2,000-1,000 RGCs/mm^2^) or degenerative (1,000-0 RGCs/mm^2^). At 10 months, AAV2.CR2-Crry-treated retinas had 31%, 61% and 8% pre-degenerative, declining, and degenerative Brn3+ RGC densities, respectively, whereas naïve retinas had 4%, 48% and 48%, and AAV2.GFP-treated retinas had 25%, 32% and 43% pre-degenerative, declining, and degenerative densities, correspondingly (**Figure 2H**). Application of Chi-squared (χ^2^) test for association indicated significant differences between AAV2.CR2-Crry compared to both naïve (*p*<0.001) and AAV2.GFP (*p*<0.01), but not in AAV2.GFP *vs.* naïve retinas (*p*=0.057; **Table 4**).

**Table 4.**
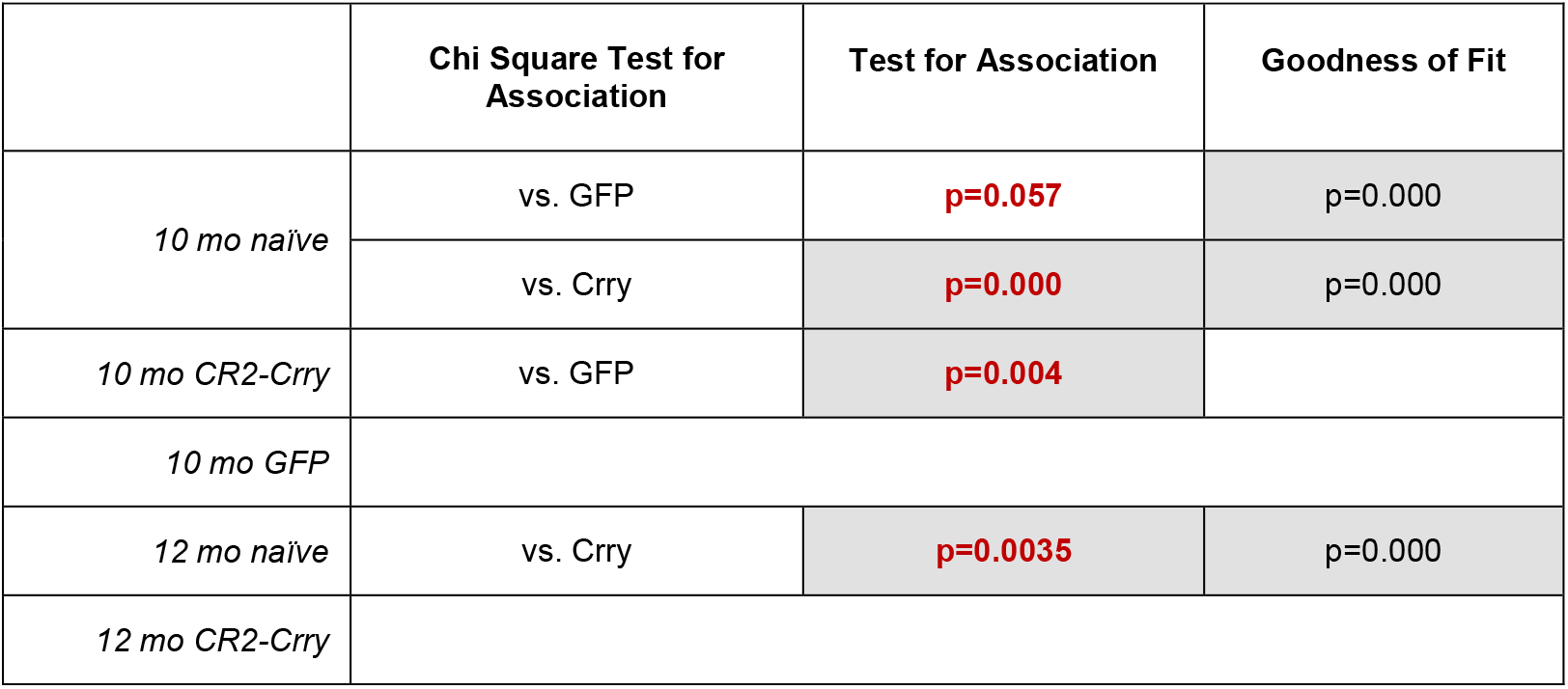
Brn3+ RGC preservation. Statistical comparison for percent retinas by level of Brn3+ RGC preservation (Figure 2H and L).

|  | Chi Square Test for Association | Test for Association | Goodness of Fit |
| --- | --- | --- | --- |
| *10 mo naïve* | vs. GFP | **p=0.057** | p=0.000 |
|  | vs. Crry | **p=0.000** | p=0.000 |
| *10 mo CR2-Crry* | vs. GFP | **p=0.004** |  |
| *10 mo GFP* |  |  |  |
| *12 mo naïve* | vs. Crry | **p=0.0035** | p=0.000 |
| *12 mo CR2-Crry* |  |  |  |

Next, to determine the magnitude of RGC depletion, we measured the extent of sectorial depletion (in degrees) of Brn3+ RGCs (less than 500 nuclei/mm^2^) and report the proportion of retinas with either no (0°), partial (1-359°) or complete depletion (360°) (**Figure 2I**). At 10 months, the AAV2.CR2-Crry group (*n*=34) had 73% of retinas with no depletion, 23% with partial and 4% with complete depletion, *vs.* 32%, 32% and 36% in naïve eyes (*n*=25), and 42%, 31% and 27% in AAV2.GFP controls (*n*=26). Chi-squared (χ^2^) test for association revealed significant differences of AAV2.CR2-Crry from either naïve (*p*<0.001) or AAV2.GFP (*p*<0.01) conditions, whereas AAV2.GFP was not significantly different from naïve retinas (p=0.522; **Table 5**). There was a significant difference across all 10 month-old groups of mean degrees of Brn3+ RGC depletion (p=0.0011, Kruskal-Wallis). Furthermore, there was a significant difference at 10 months for AAV2.CR2-Crry retinas (39°±15) compared to either naïve (160°±32, *p*<0.02) or AAV2.GFP (154°±31, *p*<0.02) (**Figure 2J**), and no significant difference between naïve and AAV2-GFP control groups by Kolmogorov-Smirnov tests (*p*=0.999; **Table 6**). This denotes a robust absence of low-density sectors in AAV2.CR2-Crry-treated retinas, indicating reduced progression towards terminal degeneration stages relative to both naïve and AAV2-GFP treated retinas. Overall, these findings show that by multiple measures AAV2.CR2-Crry results in potent preservation of Brn3+ RGCs at 10 months.

**Table 5.** RGC depletion level. Statistical comparison for percent retinas by level of Brn3+ cell depletion (Figure 2I and M).

|  | Chi Square Test for Association | Test for Association | Goodness of Fit |
| --- | --- | --- | --- |
| *10 mo naïve* | vs. GFP | **p=0.522** | p=0.286 |
|  | vs. Crry | **p=0.000** | p=0.000 |
| *10 mo CR2-Crry* | vs. GFP | **p=0.009** |  |
| *10 mo GFP* |  |  |  |
| *12 mo naïve* | vs. Crry | **p=0.000** | p=0.000 |
| *12 mo CR2-Crry* |  |  |  |

**Table 6.** RGC depletion. Statistical comparisons for mean degrees of Brn3+ cell depletion (Figure 2J and N).

|  | Normality<br>(Shapiro-Wilk) | Kruskal-Wallis |  | Dunn's Multiple<br>Comparisons test | Kolmogorov<br>-Smirnov | Mann-Whitney U |
| --- | --- | --- | --- | --- | --- | --- |
| *10 mo naïve* | p=0.000 | p=0.0011 | vs. GFP | p>0.9999 | **p=0.9992** | p=0.6788 |
|  |  |  | vs. Crry | p=0.0025 | **p=0.0114** | p=0.0005 |
| *10 mo CR2-Crry* | p=0.000 |  | vs. GFP | p=0.0126 | **p=0.0168** | p=0.0028 |
| *10 mo GFP* | p=0.000 |  |  |  |  |  |
| *12 mo naïve* | p=0.000 |  | vs. Crry | p=0.0008 | **p=0.0005** | p=0.0002 |
| *12 mo CR2-Crry* | p=0.001 |  |  |  |  |  |

Since disruption of complement pathways has been reported to have variable effects on disease progression^26, 28^, we also performed analysis at 12 months focusing on naïve retinas, which show a well-characterized progression of RGC degeneration^57, 70^, compared to AAV2-CR2-Crry treated retinas. The distributions of Brn3+ cell density showed progressive loss in both groups (**Figure 2K**). However, AAV2.CR2-Crry-treated retinas (*n*=18) retained significantly higher RGC densities compared to naïve retinas (*p*<0.01 by Kolmogorov-Smirnov, and *p*<0.001 by Mann-Whitney *U* test, *n*=19; **Table 3**). When we assessed RGC preservation at 12 months of age, we found that AAV2.CR2-Crry retinas were 11%, 50% and 39% pre-degenerative, declining and degenerative, respectively, in stark contrast to naïve retinas that were either declining or degenerative (19% and 81%, respectively) (**Figure 2L**). A Chi-squared (χ^2^) test for association indicated a significant difference between AAV2.CR2-Crry and naïve retinas (*p*<0.01; **Table 4**). Furthermore, analysis of RGC depletion at 12 months of age found that 53% AAV2.CR2-Crry retinas had no depletion, 41% had partial, and 6% had complete depletion, in contrast to 6, 19 and 75% respectively in untreated naïve retinas (**Figure 2M**). AAV2.CR2-Crry was significantly different from naïve (*p*<0.001; Chi-squared (χ^2^) test for association; **Table 5**). At 12 months of age, the mean depleted angle had increased in AAV2.CR2-Crry (108°±32) (**Figure 2N**), but was still significantly different than the untreated group (297°±31, *p*<0.001 by Kolmogorov-Smirnov; **Table 6**). The combined analysis of Brn3+ RGC maintenance and depletion demonstrates that AAV2.CR2-Crry treatment resulted in preservation of higher densities of Brn3+ RGC that were uniform across quadrants in the majority of retinas. Thus AAV2.CR2-Crry treatment results in a strong reduction in the progression towards Brn3+ RGC loss.

The neuroprotective effect of AAV2.CR2-Crry on RGC somal density may be more robust than estimated by Brn3 immunostaining, given the downregulation of this transcription factor in DBA/2J retinas at the ages analyzed^66, 67^. In line with this, AAV2.CR2-Crry treatment of a subset of 7 month-old Thy1^+/CFP^ DBA/2J reporter mice^52^ resulted in 10 month-old healthy retinas with uniformly dense mosaics of RGCs co-expressing Brn3 and Thy1 (**Figure 2O**), but also in retinas with homogeneous high densities of Thy1-expressing RGCs but loss of Brn3 expression (**Figure 2P**). Mean Thy1-positive GCL nuclear counts showed 2,031±64 in AAV2.CR2.Crry *vs.* 731±157 nuclei/mm^2^ in naïve (*n*=8 and 15, respectively), which is relatively higher than the mean density of Brn3 positive RGCs (1,565±106 nuclei/mm^2^) in AAV2.CR2-Crry retinas. This suggests transcriptional changes in persistent RGCs, and a potential underestimate of RGC somal survival by AAV2.CR2-Crry therapy if analysis is based only on Brn3 expression.

### Intraretinal axons preserve their fasciculation integrity after CR2-Crry treatment

To define whether the neuroprotective effect of CR2-Crry on RGC somata extended to the unmyelinated axon, we evaluated the integrity of axon fasciculation in the nerve fiber layer. Intraretinal RGC axons were visualized by confocal microscopy images of wholemount retinas immunostained for phosphorylated neurofilament (pNF), a cytoskeletal marker extensively used to selectively track both healthy and degenerative RGC axons^57, 66, 67, 71^. Naïve, healthy retinas displayed axons uniformly segregated in tightly packed fascicles across all quadrants in 10 month-old DBA/2J mice (**Figure 3A, A’**), whereas degenerative retinas exhibited defasciculation, with fascicle thinning and solitary axons (**Figure 3B, B’**).

**Figure 3.**
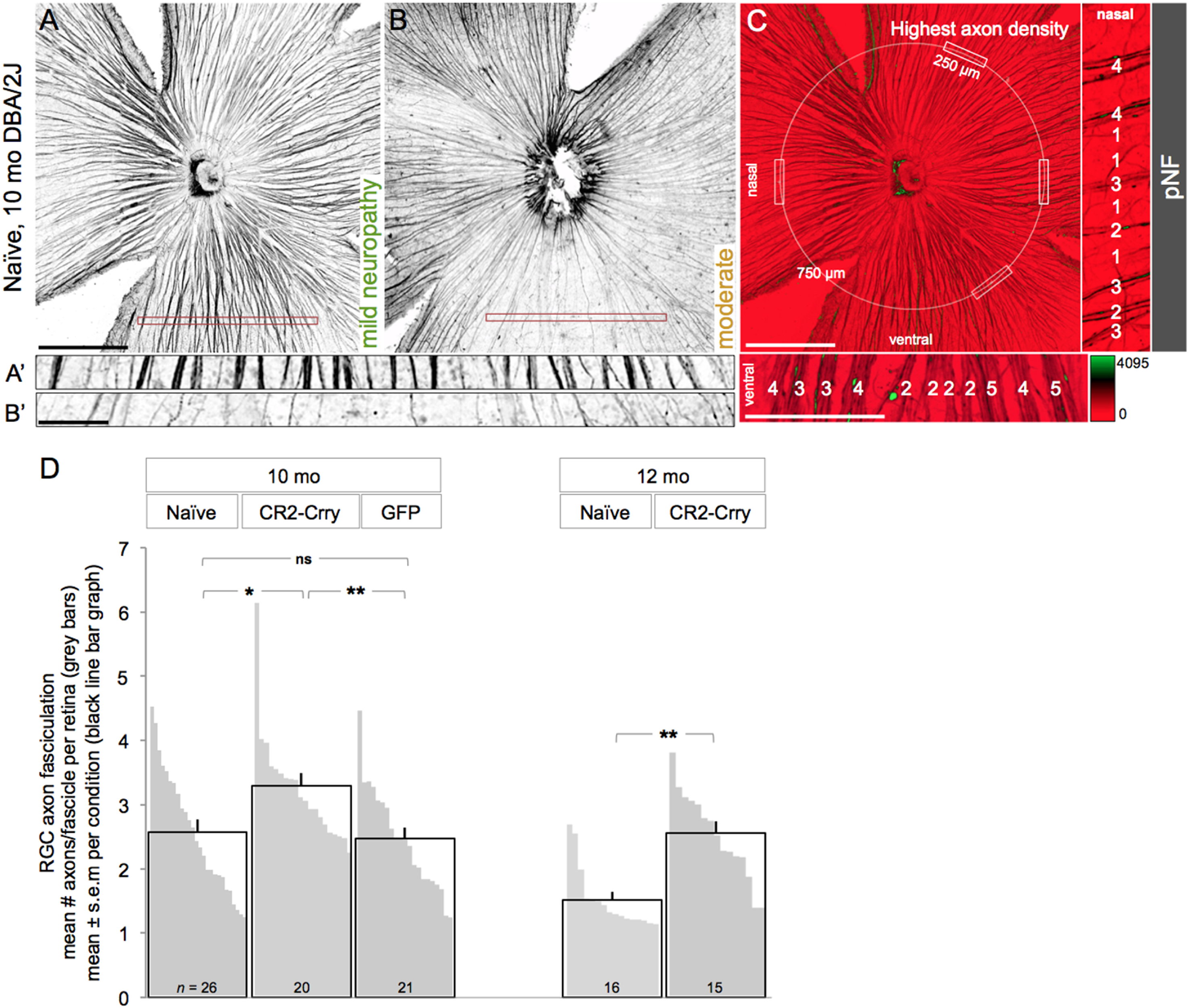
AAV2.CR2-Crry protects intraretinal axons. (**A-C**) Confocal images of pNF-labeled RGC axons on retinal wholemounts, shown as 15-μm maximum intensity projections through the NFL. Naïve, 10 month-old retinas representative of a healthy NFL throughout all quadrants (**A**), and of a NFL with sectorial degeneration, detectable by fascicle thinning and loss (**B**). Insets show high magnification views of retinal sectors with dense fascicles (A’) and a degenerated sector with sparse, single axons, near depletion (B’). The corresponding level of optic neuropathy is indicated for each retina. (**c**) Sampling method for counting axon numbers per fascicle. Axons bundled together were counted within the healthiest radius of each quadrant, along a 250-μm line, 750 μm from the ONH. Analysis was done at high magnification in pseudocolored images (scaled by fluorescence intensity), as shown in the insets (numbers of axons/fascicle indicated for samples in the nasal and ventral quadrants). Scale bars, 500 μm (A-C), and 100 μm (insets). (**D**) Number of axon per fascicle expressed as mean per experimental group (black line bars), and as mean per individual retina (grey bars). For example, the retinas shown in A and B have respectively, 3.4 and 2 axons/fascicle on average. See Table 7 for statistical analysis.

The mean density of pNF+ RGC axons per fascicle for individual retinas was estimated by counting the number of axons per bundle across 250 μm at the healthiest sector of each retinal quadrant (**Figure 3C**; see Methods for details). As a baseline, we measured 3.6±0.18 axons/fascicles (mean±s.e.m, *n*=5 retinas; data not shown) in non-glaucoma retinas from Gpnmb^wt^ DBA/2J mice at 10 months of age (**Figure S2**). At 10 months, there was a significant difference across groups (p=0.0083, One-way ANOVA; **Table 7**). AAV2.CR2-Crry-treated retinas showed significantly denser fascicles relative to naïve DBA/2J retinas (3.3±0.2 *vs.* 2.6±0.2 mean axons/fascicle for *n*=20 and 26, respectively; *p*=0.013 by an unpaired Student’s T-test) (**Figure 3D**, black line bar graphs, and **Table 7**). Retinas injected with AAV2.GFP (*n*=21) had a mean density (2.5±0.2 axons/fascicle), which was significantly reduced compared to AAV2.CR2-Crry (*p*<0.01), but not significantly different to naïve eyes (*p*=0.695; **Table 7**). Notably, AAV2.CR2-Crry retinas showed the most uniform high densities (none with less than 2.5 axons/fascicle, and more than half the retinas with more than 3 axons/fascicle). These data were confirmed by post-hoc Tukey’s multiple comparison tests (**Table 7**). To assess progression, we compared AAV2-CR2-Crry treated eyes to naïve at 12 months of age (**Figure 3D**). We observed that CR2-Crry-treated eyes showed progressive defasciculation relative to 10 months, but maintained a mean of 2.6±0.2 axons/fascicle, representing 1.7-fold more preserved axons per fascicle than age-matched naïve retinas (1.52±0.1 axons/fascicle *n*=15, *p*=0.0016 by Kolmogorov-Smirnov; **Table 7**). The significant increase in axon fasciculation resulting from AAV2.CR2-Crry, although subtle, further highlights its protective effect.

**Table 7.** Intraretinal axon fasciculation. Statistical comparisons for mean number of axons per fascicle by condition and age (Figure 3D).

|  | Normality<br>(Shapiro-Wilk) | One-way<br>ANOVA | Kruskal-Wallis |  | Tukey's multiple<br>comparison test | Unpaired<br>T-test | Kolmogorov-Smirnov |
| --- | --- | --- | --- | --- | --- | --- | --- |
| *10mo naïve* | p=0.225 | p=0.0083 | p=0.0057 | vs. GFP | p=0.9167 | **p=0.6948** | p=0.8306 |
|  |  |  |  | vs. Crry | p=0.0243 | **p=0.0129** | p=0.0091 |
| *10mo GFP* | p=0.396 |  |  | vs. Crry | p=0.0127 | **p=0.0036** | p=0.0076 |
| *10mo CR2-Crry* | p=0.001 |  |  |  |  |  |  |
| *12mo naïve* | p=0.000 |  |  | vs. Crry | p<0.0001 | p<0.0001 | **p=0.0016** |
| *12mo CR2-Crry* | p=0.873 |  |  |  |  |  |  |

### Degeneration of the optic nerve is reduced by complement inhibition

Since AAV2-CR2-Crry promotes preservation of Brn3+ RGCs and intraretinal axon fasciculation, we sought to determine whether this treatment could alter the course of axon degeneration^64, 65, 70, 72, 73^. We performed histological analysis of optic nerve sections, and individual nerves were classified as “pre-degenerative” if their proximal cross-section was dominated by dense intact axons, as “moderately degenerative” when >50% axons were healthy and 5 to 50% were dystrophic, and as “severely degenerative” when > 50% of axons were lost or dystrophic (**Figure 4A**). We then determined the ratio of nerves with pre-degenerative, moderate or severe degeneration (**Figure 4B**). In naïve untreated 7 month-old mice, the majority of nerves (96%) were pre-degenerative, and none terminally degenerative (4% moderate, and 0% severe; *n*=25), consistent with previous reports^64^. At 10 months, only half of naïve nerves remained healthy (52% pre-degenerative), and the rest showed degeneration (10% moderate, 38% severe; total *n*=83), which represents a significant change relative to 7 months (*p*<0.01 by Chi-Square; **Table 8**). In contrast, AAV2.CR2-Crry therapy from 7 to 10 months nearly eliminated nerve degeneration (90% pre-degenerative, 6% moderate, 4% severe; *n*=53), a distribution that was not significantly different from naïve 7-month-old mice by Chi-square (*p*=0.502). Relative to 10 month-old naïve nerves, AAV2.CR2.Crry treatment resulted in almost twice as many healthy nerves (90% *vs.* 52% pre-degenerative in naïve), and reduced by 10 fold the ratio of nerves at late stages of degeneration (4% *vs.* 38% severe in naïve) (*p*<0.001; **Table 8**). Age-matched AAV2.GFP eyes also showed altered incidence of nerve pathology compared to naïve, with an increased proportion of moderate and reduced proportion of severe nerves (53% mild, 29% moderate, 18% severe; *n*=34; *p*<0.05; **Table 8**). However, nerve damage in 10-month-old AAV2.GFP controls was significantly worse than 10-month-old CR2-Crry-treated nerves (*p*=0.001; **Table 8**).

**Table 8.** Statistical comparisons for proportion of optic nerve by levels of pathology (Figure 4B, C).

|  | Chi Square Test for Association | Test for Association | Goodness of Fit |
| --- | --- | --- | --- |
| *7mo naïve* | vs. 10mo naïve | **p=0.002** |  |
|  | vs. 10mo Crry | **p=0.502** |  |
|  | vs. 12mo Crry | **p=0.002** |  |
|  | vs. 15mo Crry | **p=0.016** |  |
| *10 mo naïve* | vs. 12mo naïve | **p=0.005** |  |
|  | vs. 10mo GFP | **p=0.032** | p=0.002 |
|  | vs. 10mo Crry | **p=0.000** | p=0.000 |
| *10 mo CR2-Crry* | vs. 10mo GFP | **p=0.001** |  |
|  | vs. 12mo Crry | **p=0.001** |  |
|  | vs. 15mo Crry | **p=0.058** |  |
| *10 mo GFP* | vs. 12mo GFP | **p=0.139** |  |
|  | vs. 15mo GFP | **p=0.034** |  |
| *12 mo naïve* | vs. 12mo GFP | **p=0.324** | p=0.049 |
|  | vs. 12mo Crry | **p=0.002** | p=0.000 |
| *12 mo CR2-Crry* | vs. 10mo GFP | **p=0.214** |  |
|  | vs. 15mo Crry | **p=0.974** |  |
| *12 mo GFP* | vs. 15mo GFP | **p=0.801** |  |
| *15 mo CR2-Crry* | vs. GFP | **p=0.179** |  |

**Figure 4.**
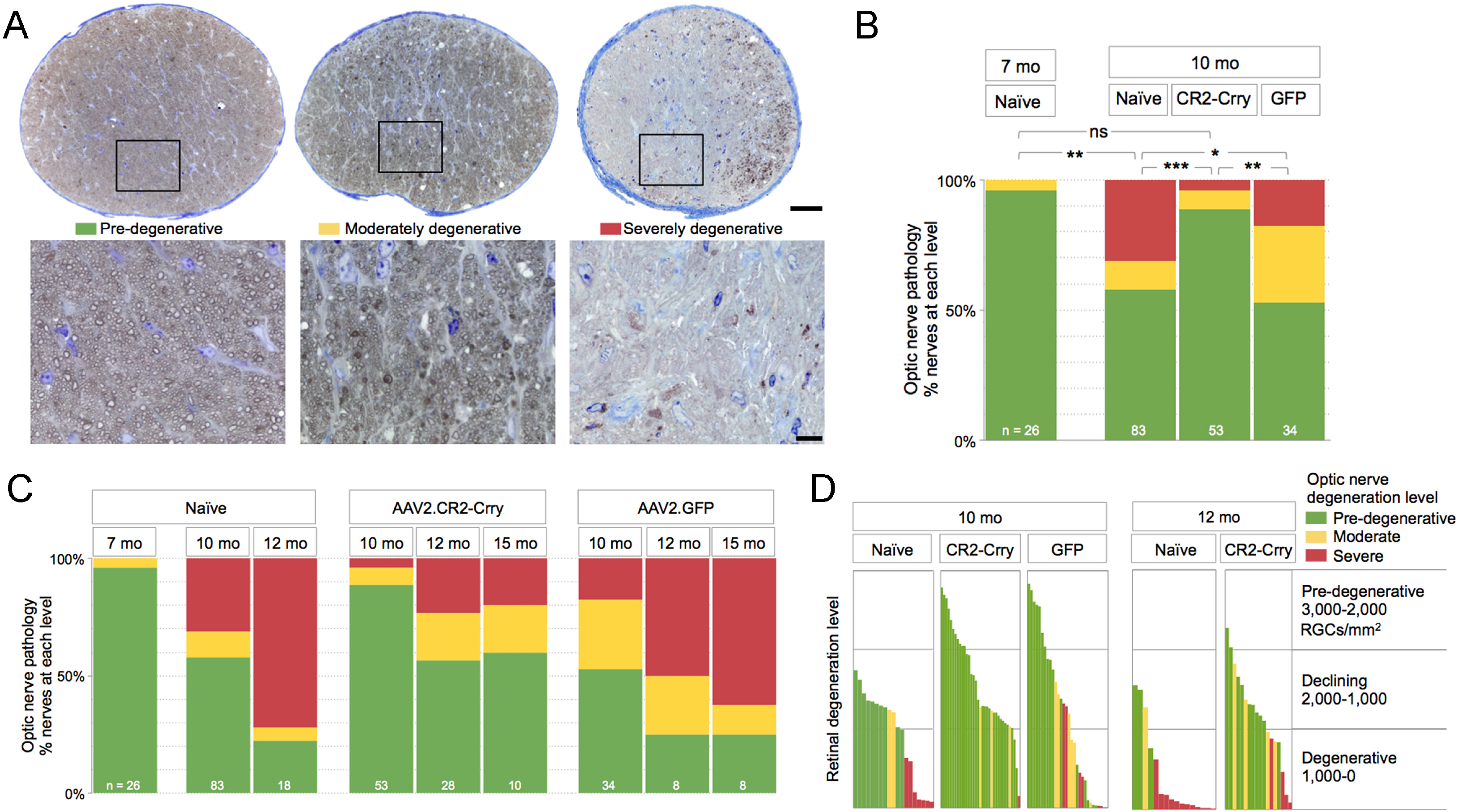
AAV2.CR2-Crry suppresses optic nerve neurodegeneration at 10 months of age, and reduces it at 12 and 15 months. (**A**) Light microscopy images of retro-orbital ON cross-sections, each representative of mild, moderate, and severe damage levels in 10 month-old naïve DBA/2J mice. Intact axons are recognizable by a clear axoplasm and intact myelin sheath, whereas dystrophic axons are detectable by a dark axoplasm and/or thick, delaminated myelin sheath. Scale bars, 50 μm (A), 10 μm (B). (**B**) Bar graph representing the distribution of ON damage by experimental group (χ^2^ test; sample size indicated). (**C**) Bar graph comparison of degeneration levels of retina *vs.* ON for individual eyes. Each bar represents the mean Brn3+ RGC density for a single retina, and its color indicates the damage level of its ON. See Table 8 for statistical analysis.

We next sought to assess progression of axon degeneration by assessing the change in ON pathology over time (**Figure 4C**). Naïve DBA/2J nerves showed progressively worsening pathology from 10 to 12 months, and by 12 months most of the nerves were severely degenerative (22% pre-degenerative, 6% moderate, 72% severe; *n*=18), relative to 10 months of age (*p*<0.01 by Chi-square; **Table 8**). Nerves from AAV2.CR2.Crry treated eyes also underwent a decline from 10 to 12 months of age (*p*=0.001), but CR2-Crry maintained significant neuroprotection at 12 months with most of the nerves healthy (62% pre-degenerative, 23% moderate, 15% severe; *n*=39; *p*<0.01; **Table 8**) relative to age-matched naïve eyes. At 15 months of age, AAV2.CR2-Crry treated nerves (*n*=10) showed continued neuroprotection, with 60% pre-degenerative, and only 20% moderately and severely degenerative each. Notably, AAV2.CR2-Crry treated eyes were not significantly more degenerative at 15 months of age than at 12 (*p*=0.974) or 10 months (*p*=0.058). In previous studies of large cohorts of naïve DBA/2J, approximately 75% of the optic nerves in females were severely damaged from 13 to 15 months of age^64^. In contrast, nerves from AAV2.GFP treated eyes showed progression of degeneration from 10 to 15 months (*p*<0.05; **Table 8**). Analysis of a limited set of AAV2.GFP nerves at 12 months (*n*=8) showed widespread damage (25% pre-degenerative, 25% and 50% moderately and severely degenerative respectively), and by 15 months AAV2.GFP nerves (*n*=8) were progressively worse with 25% pre-degenerative, 13% moderately degenerative and 63% severely degenerative (**Table 8**). Overall, these findings indicate that AAV2.CR2-Crry treatment suppressed glaucomatous ON degeneration in 10-month-old DBA/2J, and persistently delayed its progression at 12 and 15 months of age.

### AAV2.CR2-Crry treatment preserves both retina and optic nerve in individual eyes

Finally, to determine whether there is similar protection of retina and nerve by AAV2.CR2-Crry in individual eyes, we compared mean Brn3+ RGC density versus level of ON degeneration per eye (**Figure 4D**). Analysis of 10 month-old naïve eyes showed correspondence of damage levels between nerve and retina, with decline in RGC density lagging behind nerve degeneration, suggested by declining retinas with similar intermediate densities having pre-degenerative or moderately degenerative nerves. In stark contrast, age-matched AAV2.CR2.Crry-treated eyes showed preponderant preservation of pre-degenerative retinas with high to medium densities of RGC somata and matching pre-degenerative nerves. Unexpectedly, AAV2.GFP-treated eyes showed nerve protection uncoupled from retina preservation, manifest by degenerative retinas with nerves at all three stages of degeneration. At 12 months, naïve eyes showed the expected terminal degeneration of both nerve and retina, evidenced by the prevalence of nerves at severe stage with retinas depleted of RGCs. At the same age, AAV2.CR2.Crry maintained its protective effect, as suggested by the relative high RGC densities with moderately damaged nerves. This comparison underscores that AAV2.CR2-Crry ocular therapy effectively reduces progression of RGC axonal and somal decline and degeneration, protecting both the retina and its optic nerve.

## DISCUSSION

Addressing the need for treatment strategies that slow glaucoma progression, this study showed that retinal gene therapy of DBA/2J retinas with AAV2.CR2-Crry led to long-term overexpression of the Crry inhibitor and limited deposition of the C3d complement activation product on the ganglion cell layer, which strongly preserved the integrity of both ONs and retinas in most eyes, despite concurrent IOP elevation. These findings define C3 activation-dependent complement signaling as a pathological driver of glaucomatous neurodegeneration, and also establish AAV2.CR2-Crry as an effective gene therapy to locally correct pathogenic complement overactivation and restrict the onset and progression of neurodegeneration.

### Targeting the C3 activation step for complement regulation

Dysregulated or excessive complement activation underlies multiple degenerative, inflammatory and autoimmune pathologies, including neurodegenerative diseases^34, 74^. In the expanding field of complement therapeutics^75,76^, diverse approaches targeting C3 activation have been shown to be efficacious and are being extended to human disease^74, 76^, with some already in clinical trials^75^, including compstatin, a cyclic peptide that inhibits binding of C3 to the convertase^74^. Human CR1 is analogous to rodent Crry, and has been linked to CR2 to generate a targeted C3 convertase inhibitor (TT32)^77^, or modified to localize to the cell membrane (APT070)^33^. In humans C3 convertases can also be controlled via decay accelerating factor (DAF) and membrane cofactor protein (MCP)^78^, with a chimeric human CR2-DAF shown to be highly effective at C3 inhibition in mice *in vivo*^79^. Thus, our findings showing the CR2-Crry can protect RGCs from glaucomatous neurodegeneration may ultimately be translatable to human glaucoma.

### AAV-mediated gene therapy for targeted CR2-Crry delivery to the retina

*In vivo* application of systemic CR2-Crry protein has been shown to ameliorate nervous system pathology in a spectrum of animal models^40-43, 80, 81^. In these models involving trauma, neuroinflammation and ischemia, a disrupted blood-brain barrier allows access of the large CR2-Crry to the nervous system^34^. In glaucoma, complement activation is dysregulated at early stages when the retina-blood barrier is intact, which would prevent administered CR2-Crry protein from reaching C3d deposits. Retinal gene therapy based on intravitreal injection of adeno-associated virus vectors, bypasses retina-blood or other cellular barriers, and locally supplies intrinsic, long-term expression of their cargo^82^. We showed that delivery of CR2-Crry via intravitreal AAV2 provided RGCs and the inner retina with a sustained Crry overexpression that regulated pathogenic C3 overactivation. Intraretinal administration of AAV2.CR2-Crry did not increase mortality or occurrence of external infections compared to naïve mice (in over 50 treated mice; data not shown), suggesting no major systemic or off-target alterations in complement signaling, or concomitant general immunosuppression. However, it remains possible that there are extraocular effects since intravitreally delivered AAV2 has the potential to reach blood and lymphatic tissue, as demonstrated in other animal models^83^, and is able to induce innate and adaptive immune responses^84^. Locally, resident microglia are able to limit viral spread, and induce a potential neuroprotective effect secondary to this antiviral response^85^. There may also be an immune and/or inflammatory component cause by mechanical injury at the time of intravitreal injection of the viral vector^86^. Future studies will evaluate innate immune responses in AAV2-treated retinas, as well as the neuroprotective effects of their microglia, since this may mediate the modest improvement detected in eyes treated with control AAV2.GFP. Recombinant adeno-associated virus-based therapeutics is prevalent in experimental and pre-clinical ocular gene therapy, due to the unmatched versatility, efficacy and permanence of these vectors to transfer genes to the retina^87, 88^. Treatments targeted to RGCs apply intravitreal AAV2 systems optimized to selectively transduce the inner retina^47^. For translation into the clinic, AAV2 has been modified to bypass the neutralizing mechanisms particular to humans^89^, and proved safe and persistent for treating adult patients with chronic retinal disease^90^. The present study did not use AAV2-based retinal gene therapy to replace a defective RGC gene with a transgene, but to provide the glaucomatous retina with a targeted complement regulator aimed at steadily rebalancing the activation of a pathway central to the innate immune response that parallels chronic neurodegeneration.

### The complex role of local complement activation in glaucoma

Knockout studies in DBA/2J glaucoma have suggested complex roles for the complement pathway in disease progression. C1q deficiency is protective against neurodegeneration^27, 28, 30^, but C3 deficiency is detrimental, with more severe nerves at 10.5 but not 12 months of age^26^. This suggests a harmful role for C1q, and a neuroprotective role for C3^26-28^. In contrast, our study, which inhibits C3 activation but does not eliminate C3, suggests that local C3 activation contributes to RGC degeneration. This would be consistent with C1q and the classical pathway leading to C3 activation and contributing to pathology in glaucoma. It is possible that there are compensatory changes following C3 knockout that do not occur with local inhibition of C3 activation. The therapeutic value of targeted complement approaches based on tissue-specific inhibition, rather than systemic delivery or gene knockout, are supported by multiple studies^8, 33, 34^. Notably, C3 deficiency was protective against retinal ischemia-reperfusion, delaying axonal and RGC degeneration^91^, which imply a complex and more general role for complement activation following RGC injury. Therefore, when and where C1q and C3 are acting in glaucoma is still undefined. Future studies could apply inhibitors selective for each individual complement pathway to identify how each contributes to the course of glaucoma progression.

### Complement inhibition restricts onset and progression of axonal and neuronal degeneration

There is increasing evidence that degeneration of RGCs in glaucoma is compartmentalized, with distinct pathways driving axonal and somal decline programs, and with distal axonopathy preceding cell body loss in the retina^1, 64, 67, 92^. We observed that AAV2-CR2-Crry confers combined protection of both axons and somata, suggesting that complement activation mediates pathogenic events in both neuronal compartments. Onset of RGC degeneration in the DBA/2J is quite variable, with axon damage histologically detectable from 8 to 12 months of age. By applying AAV2.CR2-Crry at 7 months of age, which is generally prior to overt structural decline, we were able to constrain complement activation preceding the onset of ON degeneration, and assess effects on neurodegeneration onset and progression to terminal stage at 10 to 15 months of age. We observed highly significant preservation of ON integrity at 10, 12 and 15 months (3 to 8 months post-treatment), with a very low proportion of nerves showing severe damage. This indicates a conspicuous and continued reduction in the onset and terminal progression of pathology following complement inhibition (**Figure 5**). This protection resembles the phenotype of global C1q knockout DBA/2J mice^27, 28^, in which severe nerve pathology was eliminated at 10.5 months and reduced to < 25% at 12 months of age, suggesting delayed progression but ongoing neurodegeneration. The delay observed with AAV2-CR2-Crry is dramatic enough to offer significant potential therapeutic benefit. However, the fact that glaucoma pathology ultimately develops suggests that other pathways likely also contribute to the progression of neurodegeneration in glaucoma.

**Figure 5.**
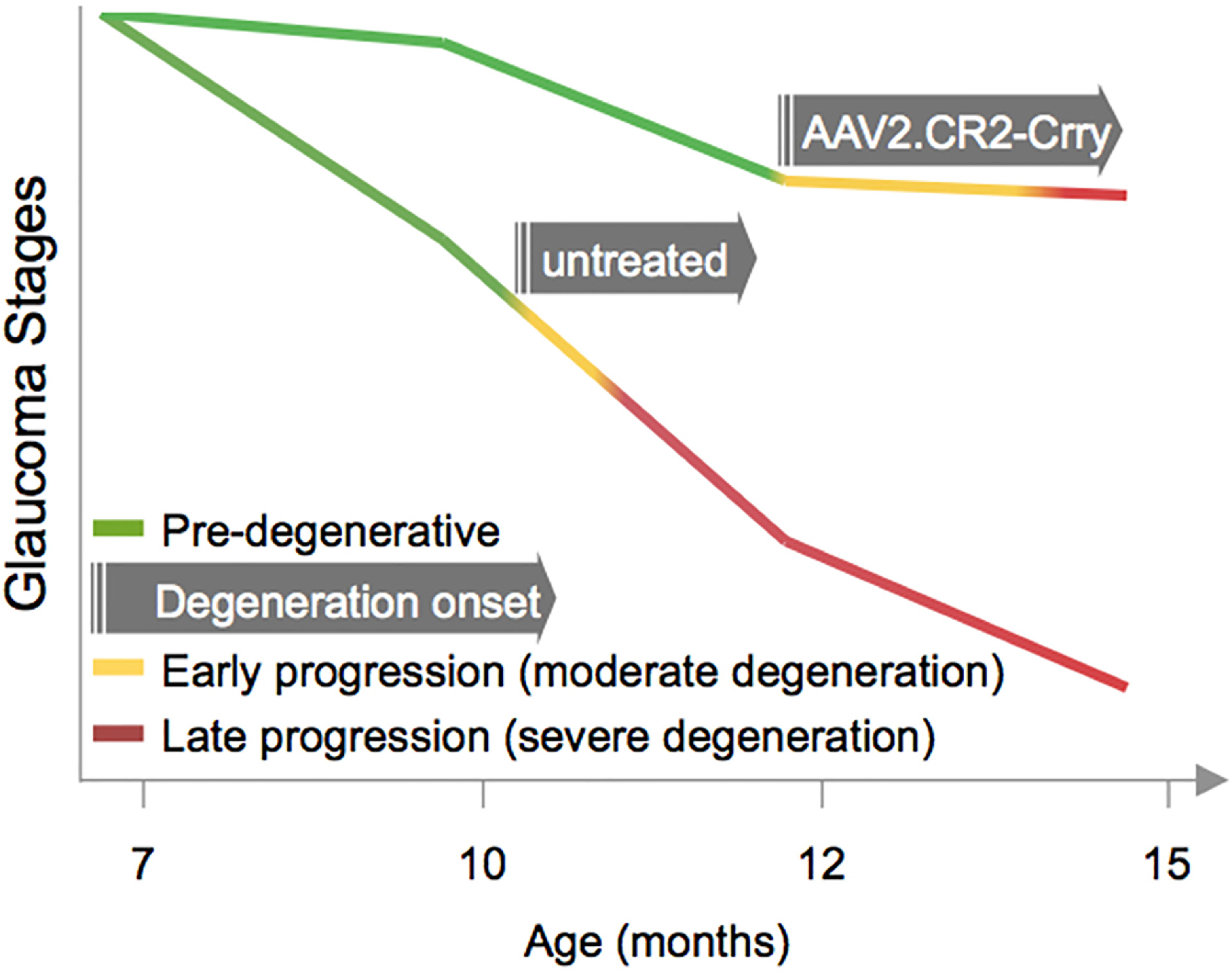
Glaucoma progression after AAV2.CR2-Crry retinal gene therapy. Diagram representing the delayed onset of ON and retinal degeneration, and slower progression in treated relative to naïve DBA/2J.

In glaucoma, the role of complement in RGC synapse remodeling and monocyte infiltration has largely been investigated in the DBA/2J model^25, 27, 28^, which is a well-characterized model of disease that exhibits many similarities to human disease, including age-dependent progression and sectorial loss^93^. However, it is possible that such immune and/or inflammatory responses are influenced by the loss of function mutation in *Gpnmb*, which is expressed in myeloid cell lineages, as recently reviewed^94^. Thus, testing AAV2.CR2-Crry gene therapy for preservation of RGC and visual function in acute models of glaucoma will be important and may reveal roles for complement during active damage *versus* chronic progression. Future molecular studies should determine the cellular sources of complement proteins or inhibitors, and define the responders, which will likely include resident microglia that express C3 receptors CR3 (CD11b/CD18, MAC-1) and infiltrating macrophages that carry recognition receptors to other complement components^10^. Recent studies suggest a dynamic capacity for microglia to sense and restrict neurodegeneration in a chronic disease such as Alzheimer’s^95^. Consistent with this notion, our previous studies in DBA/2J glaucoma revealed the spatial proximity between microglia and all neuronal compartments of RGCs^60^, and selective concentration of microglia in ONHs that were fated to develop severe neurodegeneration at advanced age^63^. How AAV2.CR2-Crry treatment modifies these innate immune responses will be the focus of future studies.

In summary, our study uncovers a damaging effect of C3 or downstream effectors in glaucoma, and establishes AAV2.CR2-Crry as a precise and translatable strategy to locally balance retinal C3 activation and reduce neurodegeneration during onset and progression of chronic glaucoma. From a clinical perspective, adeno-associated viral vector delivery of targeted complement inhibitors offer an alternative to retinal gene therapies that simply replace damaged genes, by therapeutically regulating pathogenic complement pathways in chronic glaucoma, an approach that is applicable to other CNS neurodegenerative diseases.

## MATERIALS AND METHODS

### Mice

Experimental work, handling and care of mice were carried out in compliance with the ARVO Statement for the Use of Animals in Ophthalmic and Vision Research, and under an animal protocol approved by the University of Utah Institutional Animal Care and Use Committee. DBA/2J and non-glaucoma control Gpnmb^+/+^ DBA/2J mice were obtained from Jackson Laboratories (Bar Harbor, ME) or bred in-house under sterile conditions, with 12/12 diurnal light cycles and food and water *ad libitum*. This study used females only, and each experimental group included similar numbers of mice born in Utah and Maine. Transgenic Thy1^CFP/+^ DBA/2J mice, which express cyan fluorescent protein reporter under the Thy1 locus, were maintained by crossing to DBA/2J, using the original genotype to identify heterozygous offspring^52^.

### AAV2-vector production

Recombinant, self-complementary adeno-associated virus (scAAV2) with triple-mutant capsids (Y444+500+730F) were generated by site-directed mutagenesis of surface tyrosine residues on the VP3 protein, and prepared as previously described^47, 54^. The titer of DNase-resistant scAAV2 vector genomes was measured by real-time PCR relative to a standard; purity was validated by silver-stained sodium dodecyl sulfate–polyacrylamide gel electrophoresis, sterility and absence of endotoxin were assayed, aliquots were stored at −80°C, and thawed no more than twice before use.

### Intravitreal injection

Mice were anesthetized by intraperitoneal injection of Avertin (1.3% 2,2,2-tribromoethanol and 0.8% tert-amyl alcohol, Sigma-Aldrich, St Louis, MO), and their eyes with topical tetracaine hydrochloride ophthalmic solution (Alcon, Forth Worth, TX). A puncture through the wall of the eye was made at the posterior angle of the eye, below the limbus, with a 31G needle held by hand and observed under a dissecting microscope with fiber optic light. Immediately after, 2 μL of AAV2 vector particles (10^10^ total vector genomes) were dispensed through this aperture using a blunt 34G needle attached to a 5-μl Nanofil Hamilton syringe ran by an Ultra Micro Pump with Micro-4 controller (World Precision Instruments, Sarasota, FL). After delivery of the vector at a rate of 1μl/min the needle was kept inside the eye for 60 sec. Finally, the eyes were covered with erythromycin ophthalmic ointment (Bausch+Lomb, Rochester, NY), and the animal was kept warm over a warm-water blanket until recovering full motility. Individual mice were injected with AAV2 expressing either the coding sequence of CR2-Crry or green fluorescent protein (GFP) from the chicken β-actin promoter (CBA).

### Intraocular pressure (IOP)

IOP was determined as previously described^96^. Briefly, mice were sedated by inhalation of isoflurane via a Rodent Circuit Controller (VetEquip, Inc., Pleasanton, CA), delivering 2% isoflurane in 2 L/min oxygen within a sealed box until lack of motility, and through a nose cone afterwards. Within 3 minutes of the start of anesthesia, 12 separate IOP measurements were performed using a Tonolab (Icare Tonovet, Finland), of which 10 were averaged (excluding the lowest and highest for each measure). Naïve and treated mice had their IOP measured monthly from 7 to 10 months of age.

### Tissue collection and fixation

Mouse eyes were collected as previously described^68^. Briefly, full anesthesia was induced by inhalation of 2% (vol) Isoflurane in 2 liter/min oxygen, then mice were transcardially perfused with 5 ml PBS followed by 20 ml 4% paraformaldehyde (PFA; Electron Microscopy Sciences, Hatfield, PA) in 0.1 M PBS, circulated with a peristaltic pump (Dynamax, Rainin, Oakland, CA). Eyes and ONs were exposed *in situ* and post-fixed for 2 h in PFA at 4°C, then nerves were dissected and fixed overnight in 1.2% PFA and 0.8% glutaraldehyde in phosphate buffer (Electron Microscopy Sciences; Hatfield, PA).

### Whole-retina qRT-PCR

Retinas were prepared and processed for RNA isolation not exceeding 10 minutes, as previously described^68^. Briefly, each individual retina was dissected fresh under RNAse-free conditions, mechanically dissociated. As recently described^96^, RNA was isolated using the RNeasy micro kit (Qiagen, Germantown, MD), and its integrity determine by Bioanalyzer (discarded < 8) (Agilent, Salt Lake City, UT). We used Superscript III First Strand cDNA synthesis (Invitrogen, Carlsbad CA) and Nano-drop spectrophotometry (ThermoFisher Scientific, Waltham, MA). Quantitative real-time PCR reactions were prepared using SYBR Select Master mix (ThermoFisher Scientific), and run on the 7900 HT Fast Real-Time PCR system with QuantStudio 12K Flex software (Applied Biosystems, Foster City, CA). qPCR analysis was performed with the ΔΔC_t_ method to determine relative expression changes^97^, and gene expression was normalized to 3 housekeeping genes, including Gapdh, as well as Rik, and Mrps6 that have consistent expression in the DBA/2J retina^27^. Primers were designed using Primer-BLAST software^98^ as follows (forward primer fist, reverse primer second, in 5’ to 3’ orientation), *Gapdh*: TGCACCACCAACTGCTTAGC, GGCATGGACTGTGGTCATGAG; *Cpsf7*/*5730453I16Rik*: TGTCCCTCCTCCTCCTCCTG, GGGGGAGGTACAGCCAGATG; *Mrps6*: AATCCCTGATGGACCGAGGA, TGTGCTGCTGGCTGTGACTC; *Crry*: CCAGCAGTGTGCATTGTCAGTCC, CCCCTTCTGGAATCCACTCATCTC^99^.

### Optic nerve histology, imaging and analysis

Nerves were prepared as previously described^73^. Briefly, the postlaminar 1-1.5 mm of myelinated nerve was processed as 1-μm thick plastic cross-sections that were stained with toluidine-blue to increase contrast for light microscopy, and with paraphenylenediamine (PPD) to identify dystrophic myelin and axoplasm^64^. High-resolution multipoint (36 xy) images were generated using compound light-microscopy and a 60x lens (BX51 and cellSens software, Olympus, Center Valley, PA). Three authors, using images masked for corresponding experimental group and retinal pathology, independently evaluated ON damage.

### Retinal histology and immunofluorescence

Retinas were prepared as wholemounts the day after perfusion, and immunostained as previously described the following day^68, 100, 101^. Briefly, wholemount retinas were incubated with primary antibodies for 3 days at 4°C, then with secondary antibodies for 2 hr. Primary antibodies included: mouse anti-phosphorylated neurofilament, pNF, (M07062 Dako, Carpinteria, CA; diluted 1:100), mouse anti-non-phosphorylated neurofilament H, NF-H (801703 Biolegend, San Diego CA; diluted 1:500), goat anti-Brn3 (C-13, sc-6026 Santa Cruz Biotechnology, Santa Cruz CA; diluted 1:50), and goat anti-C3d^59, 60^ (AF2655 R&D Systems, Minneapolis, MN; diluted 1:1:500). Secondary antibodies included donkey anti-IgG Alexa-Fluor 488, 555 and 647 nm (Invitrogen, La Jolla, CA; diluted 1:400).

### Confocal imaging

Confocal images spanning entire retinal flat mounts were generated as previously described^68, 100^, using a confocal imaging system equipped with a 20× lens and a resonant scanner (A1R confocal, Eclipse Ti inverted microscope and NIS-Elements C, Nikon). Multipoint images (625 xy positions) were acquired at high resolution (0.41 μm/px), then stitched and projected as maximal intensity of the inner 30-40 μm of retina (0.8-μm step). To allow image analysis and quantification, the parameters of image acquisition were maintained constant between retinal samples, and, for illustration, images had their brightness and contrast minimally edited.

### Brn3-expressing cell quantification

To estimate “RGC preservation”, we visually identified in each retinal quadrant the sector with densest Brn3+ nuclei at mid-periphery, where the number of Brn3+ nuclei was counted within a 250×250 μm box, masked to condition and age. The average of the four quadrants represents the mean density of preserved (highest) Brn3+ cells per retina, based on which individual retinas were considered as pre-degenerative (≥ 2,000 cells/mm^2^), declining (2,000-1,000 cells/mm^2^), or degenerative (≤ 1,000 cells/mm^2^). To estimate “RGC depletion”, we measured the total extent of retinal sectors (in degrees of an angle with the apex at the ONH) with ≤ 500 Brn3+ nuclei/mm^2^), classifying individual retinas as having no (0°), partial (1-359°) or complete depletion (360°) of Brn3+ RGCs.

### Intraretinal axon fasciculation quantification

The density of RGC axon fasciculation was analyzed retinal wholemounts immunostained for phosphorylated neurofilament (pNF). Confocal images were projected at maximal intensity to span the entire nerve fiber layer (NFL), and were pseudocolored and zoomed 400x to facilitate the visualization of single axons. First, retinal sectors with preserved axon fasciculation were visually selected in each retinal quadrant, and a 250 μm-long line was traced perpendicular to the fascicles at an eccentricity of 750 μm (**Figure 3C**). Blind to experimental group, the number of pNF+ axons along was counted each of the four lines. Counts included axons bundled together in a fascicle, as well as defasciculated, single axons.

### Statistical analysis

All data were analyzed using statistical software (SPSS Statistics 23, IBM, Armonk, NY) or GraphPad Prism (La Jolla, CA). First, normality was determined by the Shapiro-Wilk test. Depending on the results of normality tests, either a One-Way ANOVA or a Kruskal-Wallis test was used to determine differences across groups. Then a follow-up post-hoc multiple comparison test was applied: Dunn’s multiple comparisons for non-normal data or Tukey’s for normal data. Normal data sets were further tested using Students Unpaired T-test (with a Welch’s correction if the variances between the two were significantly different). Non-normal data sets (Brn3+ cell density, mean degrees of Brn3+ RGC depletion, mean axon fascicles at 12 months, and IOP) were then further compared by two different non-parametric statistical tests, Kolmogorov-Smirnov and Mann-Whitney U. Differences in Brn3+ RGC preservation and depletion, as well as level of optic nerve degeneration were determined by a Pearson Chi-square (χ^2^) test for association. χ^2^ Goodness of Fit test was run and supplied in the supplementary data. We used a 95% confidence interval, and a *p* value of < 0.05 was set for rejecting the null hypothesis.

## ACKNOWLEDGEMENTS

The authors acknowledge research support to M.L.V. from the National Eye Institute 1R01EY023621 and 1R01EY020878, the Glaucoma Research Foundation and Melza M. and Frank Theodore Barr Foundation, to W.W.H. from the Macula Vision Research Foundation, and Research to Prevent Blindness, Inc., and to S.T. from the Dept. of Veteran’s Affairs RX001141 and BX001201. The authors thank Dr. Nicholas Brecha (University of California Los Angeles) for founder Thy1^CFP/+^ DBA/2J mice. WWH and the University of Florida have a financial interest in the use of AAV therapies, and own equity in a company (AGTC, Alachua, Florida) that might, in the future, commercialize some aspects of this work. ST is an inventor on a licensed patent that covers CR2-targeted complement inhibitors, and has received royalties from Alexion (New Haven, CT).

## AUTHOR CONTRIBUTIONS

Conceptualization and Methodology, A.B. and M.L.V.; Investigation, A.B., S.R.A., K.T.B., C.O.R., M.R.S., V.A.C., and S.L.B.; Visualization, Writing – Original Draft, A.B.; Writing – Review & Editing, A.B., M.L.V., W.W.H., and S.T.; Funding Acquisition, M.L.V., W.W.H., and S.T.; Resources, M.L.V., W.W.H., and S.T.; Project Administration, A.B.

## SUPPLEMENTAL FIGURES

**Figure S1**. C3d immunostaining specificity. (**A**) Confocal image of the GCL/NFL (30 μm-maximal intensity projection) from a naïve Thy1^CFP^ DBA/2J retinal wholemount triple-immunostained for C3d, SMI32 and gamma-synuclein (same retina shown in Figure 1F). Arrows point to RGCs showing C3d deposition and co-immunostaining for Thy1, SMI32 and/or ϒ-synuclein. (**B**) Confocal image of a retinal wholemount triple-immunostained for C3d, SMI32 and Iba1 (0.3 μm single optical slice). Asterisks indicate C3d-expressing SMI31+ α-RGCs. Crosses indicate C3d-negative Iba1+ microglia.

**Figure S2**. Intraretinal axon fasciculation (**A**). Confocal image of a retinal wholemount from a non-glaucoma Gpnmb^WT^ DBA/2Jmouse, immunostained for pNF (maximal intensity projection of 20 μm). Scale, 500 μm.

